## Supplementary Tables for "Antibodies utilizing VL6-57 light chains target a convergent cryptic epitope on SARS-CoV-2 spike protein driving the genesis of Omicron variants"

**Supplementary Table 1 |** Summary of rate constants (kon, koff) and dissociation
constants (K_D_) for the R1-26 interactions in Fig. 1c,d

| **Ligand** | **Analyte** | ***k*_on_ (M^−1^ s^−1^)** | ***k*_off_ (s^−1^)** | ***K_D_* (nM)** |
| --- | --- | --- | --- | --- |
| R1-26 | RBD (WT) | 4.60×10^5^ | 1.77×10^-3^ | 3.84 |
|  | RBD (Alpha) | 6.25×10^5^ | 6.90×10^-3^ | 11.05 |
|  | RBD (Beta) | 5.39×10^5^ | 5.27×10^-3^ | 9.78 |
|  | RBD (Delta) | 3.41×10^5^ | 3.51×10^-3^ | 10.29 |
|  | RBD (Omicron BA.1) | - | - | no binding |
|  | S-trimer (WT) | 3.80×10^3^ (*k*_on1_) | 1.37×10^-6^ (*k*_off1_) | 0.36 (*k*_off1_/*k*_on1_) |
|  |  | 2.38×10^4^ (*k*_on2_) | ＜1.00×10^-7^ (*k*_off2_) | ≤0.058 (*k*_off1_/*k*_on2_) |
|  | S-trimer (Alpha) | 1.08×10^4^ (*k*_on1_) | 3.27×10^-5^ (*k*_off1_) | 3.04 (*k*_off1_/*k*_on1_) |
|  |  | 6.65×10^4^ (*k*_on2_) | 2.29×10^-7^ (*k*_off2_) | 0.49 (*k*_off1_/*k*_on2_) |
|  |  |  |  | ≤0.021 (*k*_off2_/*k*_on1_) |
|  | S-trimer (Beta) | 1.21×10^4^ (*k*_on1_) | 1.48×10^-5^ (*k*_off1_) | 1.22 (*k*_off1_/*k*_on1_) |
|  |  | 2.69×10^4^ (*k*_on2_) | ＜1.00×10^-7^ (*k*_off2_) | 0.55 (*k*_off1_/*k*_on2_) |
|  |  |  |  | ＜0.0083 (*k*_off2_/*k*_on1_) |
|  | S-trimer (Delta) | 1.71×10^4^ (*k*_on1_) | 2.95×10^-6^ (*k*_off1_) | 0.17 (*k*_off1_/*k*_on1_) |
|  |  | 2.47×10^4^ (*k*_on2_) | ＜1.00×10^-7^ (*k*_off2_) | 0.12 (*k*_off1_/*k*_on2_) |
|  |  |  |  | ＜0.0059 (*k*_off2_/*k*_on1_) |
|  | S-trimer (Omicron BA.1) | 5.29×10^3^ (*k*_on1_) | 2.54×10^-4^ (*k*_off1_) | 47.99 (*k*_off1_/*k*_on1_) |
|  |  | 9.09×10^3^ (*k*_on2_) | 1.46×10^-4^ (*k*_off2_) | 27.94 (*k*_off1_/*k*_on2_) |
|  |  |  |  | 27.64 (*k*_off2_/*k*_on1_) |
|  |  |  |  | 16.09 (*k*_off2_/*k*_on2_) |

**Supplementary Table 2 | Cryo-EM data collection, refinement and validation statistics of S-GSAS/6P:R1-26, S-GSAS/6P:H18 and** **S-GSAS/6P:H18:R1-32**

|  | S-GSAS/6P: R1-26   3:2 | S-GSAS/6P: R1-26  3:3 | S-GSAS/6P: R1-26  6:6 C1 symmetry | RBD:R1-26 /Focused | S-GSAS/6P: H18  3:2 |
| --- | --- | --- | --- | --- | --- |
| **Data collection and processing** |  |  |  |  |  |
| Magnification | 81,000 | 81,000 | 81,000 | 81,000 | 45,000 |
| Voltage (kV) | 300 | 300 | 300 | 300 | 200 |
| Electron exposure (e^–^/Å^2^) | 50 | 50 | 50 | 50 | 60 |
| Defocus range (μm) | 0.8-2.0 | 0.8-2.0 | 0.8-2.0 | 0.8-2.0 | 0.8-2.5 |
| Pixel size (Å) | 1.095 | 1.095 | 1.095 | 1.095 | 0.88 |
| Movies (no.) | 4,343 | 4,343 | 4,343 | 4,343 | 4,822 |
| Initial particle images (no.) | 2,359,986 | 2,359,986 | 2,359,986 | 2,359,986 | 2,357,740 |
| Symmetry imposed | *C1* | *C1* | *C1* | *C1* | *C1* |
| Final particle images (no.) | 110,522 | 388,973 | 43,870 | 1,166,919 | 42,170 |
| Map resolution (Å) | 3.44 | 3.19 | 5.31 | 3.54 | 5.56 |
| FSC threshold | 0.143 | 0.143 | 0.143 | 0.143 | 0.143 |
| Map resolution range (Å) | 3.16-11.51 | 2.99-10.63 | 4.09-19.19 | 3.38-5.72 | 4.40-19.60 |
| **Refinement** |  |  |  |  |  |
| Initial model used | PDB 7YEG | PDB 7YEG | PDB 7YEG | PDB 7YEG | PDB 7YEG |
| Model resolution (Å) | 3.8 | 3.4 | 7.2 | 3.8 | 7.1 |
| FSC threshold | 0.5 | 0.5 | 0.5 | 0.5 | 0.5 |
| Map sharpening *B* factor (Å^2^) |  |  |  | -112 |  |
| Model composition |  |  |  |  |  |
| Non-hydrogen atoms | 32,311 | 35,558 | 70,965 | 3,310 | 32,374 |
| Protein residues | 4,068 | 4,503 | 8,985 | 425 | 4,072 |
| Ligands | 60 | 60 | 120 | 2 | 57 |
| *B* factors (Å^2^) |  |  |  |  |  |
| Protein | 362.62 | 301.53 | 367.59 | 143.06 | 526.83 |
| Ligand | 248.51 | 194.03 | 311.95 | 141.06 | 326.68 |
| R.m.s. deviations |  |  |  |  |  |
| Bond lengths (Å) | 0.003 | 0.003 | 0.003 | 0.004 | 0.002 |
| Bond angles (°) | 0.633 | 0.611 | 0.609 | 0.758 | 0.548 |
| **Validation** |  |  |  |  |  |
| MolProbity score | 1.60 | 1.61 | 1.62 | 1.60 | 1.53 |
| Clashscore | 5.76 | 5.98 | 5.79 | 5.13 | 5.85 |
| Poor rotamers (%) | 0.60 | 0.70 | 0.37 | 0.84 | 0.08 |
| Ramachandran plot |  |  |  |  |  |
| Favored (%) | 95.82 | 95.91 | 95.59 | 95.23 | 96.67 |
| Allowed (%) | 4.18 | 4.09 | 4.41 | 4.77 | 3.33 |
| Disallowed (%) | 0.00 | 0.00 | 0.00 | 0.00 | 0.00 |

|  | S-GSAS/6P: H18  3:3 | S-GSAS/6P: H18  6:6 C1 symmetry | S-GSAS/6P: H18  6:6 C3 symmetry | S-GSAS/6P: H18:R1-32  3:2:2 | S-GSAS/6P: H18:R1-32  3:3:3 |
| --- | --- | --- | --- | --- | --- |
| **Data collection and processing** |  |  |  |  |  |
| Magnification | 45,000 | 45,000 | 45,000 | 130,000 | 130,000 |
| Voltage (kV) | 200 | 200 | 200 | 300 | 300 |
| Electron exposure (e^–^/Å^2^) | 60 | 60 | 60 | 50 | 50 |
| Defocus range (μm) | 0.8-2.5 | 0.8-2.5 | 0.8-2.5 | 0.6-2.0 | 0.6-2.0 |
| Pixel size (Å) | 0.88 | 0.88 | 0.88 | 0.93 | 0.93 |
| Movies (no.) | 4,822 | 4,822 | 4,822 | 9,861 | 9,861 |
| Initial particle images (no.) | 2,357,740 | 2,357,740 | 2,357,740 | 3,335,676 | 3,335,676 |
| Symmetry imposed | *C1* | *C1* | *C3* | *C1* | *C1* |
| Final particle images (no.) | 32,427 | 10,076 | 16,440 | 19,648 | 140,082 |
| Map resolution (Å) | 6.12 | 8.38 | 5.80 | 3.96 | 3.42 |
| FSC threshold | 0.143 | 0.143 | 0.143 | 0.143 | 0.143 |
| Map resolution range (Å) | 4.62-19.17 | 6.69-24.88 | 4.49-19.01 | 3.58-19.47 | 3.18-12.85 |
| **Refinement** |  |  |  |  |  |
| Initial model used | PDB 7YEG | PDB 7YEG | PDB 7YEG | PDB 7YEG | PDB 7YEG |
| Model resolution (Å) | 6.9 | 10.9 | 7.7 | 4.0 | 3.6 |
| FSC threshold | 0.5 | 0.5 | 0.5 | 0.5 | 0.5 |
| Map sharpening *B* factor (Å^2^) |  |  |  |  |  |
| Model composition |  |  |  |  |  |
| Non-hydrogen atoms | 35,629 | 70,986 | 71,271 | 38,806 | 45,309 |
| Protein residues | 4,508 | 8,980 | 9,018 | 4,950 | 5,829 |
| Ligands | 57 | 114 | 114 | 57 | 57 |
| *B* factors (Å^2^) |  |  |  |  |  |
| Protein | 312.67 | 538.26 | 349.39 | 340.88 | 390.49 |
| Ligand | 190.03 | 503.00 | 335.34 | 190.03 | 190.03 |
| R.m.s. deviations |  |  |  |  |  |
| Bond lengths (Å) | 0.003 | 0.002 | 0.003 | 0.004 | 0.002 |
| Bond angles (°) | 0.546 | 0.542 | 0.611 | 0.967 | 0.537 |
| **Validation** |  |  |  |  |  |
| MolProbity score | 1.62 | 1.52 | 1.60 | 1.62 | 1.60 |
| Clashscore | 6.42 | 5.38 | 6.29 | 6.68 | 6.29 |
| Poor rotamers (%) | 0.71 | 0.09 | 0.20 | 0.59 | 0.76 |
| Ramachandran plot |  |  |  |  |  |
| Favored (%) | 96.12 | 96.48 | 96.20 | 96.32 | 96.29 |
| Allowed (%) | 3.88 | 3.52 | 3.80 | 3.68 | 3.71 |
| Disallowed (%) | 0.00 | 0.00 | 0.00 | 0.00 | 0.00 |

|  | S-GSAS/6P: H18:R1-32  3:3:3 one RBD rotated | S1:H18:R1-32  1:1:1 | S1:H18:R1-32  2:2:2 |
| --- | --- | --- | --- |
| **Data collection and processing** |  |  |  |
| Magnification | 130,000 | 130,000 | 130,000 |
| Voltage (kV) | 300 | 300 | 300 |
| Electron exposure (e^–^/Å^2^) | 50 | 50 | 50 |
| Defocus range (μm) | 0.6-2.0 | 0.6-2.0 | 0.6-2.0 |
| Pixel size (Å) | 0.93 | 0.93 | 0.93 |
| Movies (no.) | 9,861 | 9,861 | 9,861 |
| Initial particle images (no.) | 3,335,676 | 3,335,676 | 3,335,676 |
| Symmetry imposed | *C1* | *C1* | *C2* |
| Final particle images (no.) | 13,185 | 178,003 | 144,598 |
| Map resolution (Å) | 4.13 | 3.57 | 3.68 |
| FSC threshold | 0.143 | 0.143 | 0.143 |
| Map resolution range (Å) | 3.67-21.44 | 3.44-10.00 | 3.53-16.12 |
| **Refinement** |  |  |  |
| Initial model used | PDB 7YEG | PDB 8HC5 | PDB 8HC5 |
| Model resolution (Å) | 4.4 | 3.9 | 4.0 |
| FSC threshold | 0.5 | 0.5 | 0.5 |
| Map sharpening *B* factor (Å^2^) |  | -74 | -75.5 |
| Model composition |  |  |  |
| Non-hydrogen atoms | 45,309 | 11,429 | 22,856 |
| Protein residues | 5,829 | 1,498 | 2,998 |
| Ligands | 57 | 1 | 2 |
| *B* factors (Å^2^) |  |  |  |
| Protein | 162.70 | 195.18 | 182.59 |
| Ligand | 134.02 | 74.48 | 37.02 |
| R.m.s. deviations |  |  |  |
| Bond lengths (Å) | 0.003 | 0.002 | 0.002 |
| Bond angles (°) | 0.628 | 0.518 | 0.525 |
| **Validation** |  |  |  |
| MolProbity score | 1.63 | 1.42 | 1.55 |
| Clashscore | 6.37 | 4.00 | 5.80 |
| Poor rotamers (%) | 0.46 | 0.31 | 0.04 |
| Ramachandran plot |  |  |  |
| Favored (%) | 95.95 | 96.49 | 96.49 |
| Allowed (%) | 4.05 | 3.51 | 3.51 |
| Disallowed (%) | 0.00 | 0.00 | 0.00 |

Supplementary Table 3 | Kinetic parameters of the recombinational mAbs binding to RBDs of SARS-CoV-2, SARS-CoV-1, Pangolin CoV GD1 and Bat CoV RaTG13 and in Figs. 5a,b and S11

|  | **SARS-CoV-2** | | | **SARS-CoV-1** | | | **Pangolin CoV GD1** | | | **Bat CoV RaTG13** | | |
| --- | --- | --- | --- | --- | --- | --- | --- | --- | --- | --- | --- | --- |
|  | *k*_on_ (M^−1^ s^−1^) | *k*_off_ (s^−1^) | *K_D_* (nM) | *k*_on_ (M^−1^ s^−1^) | *k*_off_ (s^−1^) | *K_D_* (nM) | *k*_on_ (M^−1^ s^−1^) | *k*_off_ (s^−1^) | *K_D_* (nM) | *k*_on_ (M^−1^ s^−1^) | *k*_off_ (s^−1^) | *K_D_* (nM) |
| **H4** | 3.51×10^5^ | 1.05×10^-2^ | 29.95 | - | - | no binding | 2.88×10^4^ | 1.02×10^-2^ | 355 | 2.67×10^4^ | 1.34×10^-2^ | 503 |
| **H5** | 1.29×10^5^ | 1.95×10^-1^ | 1504 | - | - | no binding | - | - | no binding | - | - | no binding |
| **H14** | 5.36×10^5^ | 1.82×10^-2^ | 34.05 | - | - | no binding | 4.55×10^4^ | 2.05×10^-2^ | 451 | 4.99×10^4^ | 1.51×10^-2^ | 302 |
| **H16** | 5.09×10^5^ | 4.98×10^-2^ | 97.9 | - | - | no binding | 3.44×10^4^ | 2.54×10^-2^ | 737 | - | - | no binding |
| **H18** | 3.16×10^5^ | 9.31×10^-3^ | 29.49 | 1.23×10^4^ | 2.83×10^-3^ | 231 | 4.65×10^4^ | 6.24×10^-3^ | 134 | 1.84×10^4^ | 6.12×10^-3^ | 332 |

Supplementary Table 4 | Kinetic parameters of R1-26, H18 and S2A4 binding to the generated RBD mutants in Fig. 6c

|  | **R1-26** | | | **H18** | | | **S2A4** | | |
| --- | --- | --- | --- | --- | --- | --- | --- | --- | --- |
|  | *k*_on_ (M^−1^ s^−1^) | *k*_off_ (s^−1^) | *K_D_* (nM) | *k*_on_ (M^−1^ s^−1^) | *k*_off_ (s^−1^) | *K_D_* (nM) | *k*_on_ (M^−1^ s^−1^) | *k*_off_ (s^−1^) | *K_D_* (nM) |
| **Wuhan-Hu-1** | 5.07×10^5^ | 2.91×10^-3^ | 5.75 | 5.60×10^5^ | 1.62×10^-2^ | 28.99 | 3.80×10^5^ | 9.60×10^-3^ | 25.27 |
| **Omicron** | - | - | no binding | - | - | no binding | - | - | no binding |
| **Omicron-L371S** | - | - | no binding | - | - | no binding | - | - | no binding |
| **Omicron-P373S** | - | - | weak binding | - | - | no binding | - | - | weak binding |
| **Omicron-F375S** | - | - | no binding | - | - | no binding | - | - | no binding |
| **Omicron-(L371S+P373S)** | 1.36×10^5^ | 2.31×10^-3^ | 16.94 | 1.71×10^5^ | 2.84×10^-2^ | 166 | 1.07×10^5^ | 9.58×10^-3^ | 89.39 |
| **Omicron-(P373S+F375S)** | 1.59×10^6^ | 7.85×10^-2^ | 49.34 | 1.31×10^6^ | 7.78×10^-2^ | 59.56 | 7.07×10^5^ | 2.73×10^-2^ | 38.62 |
| **Omicron-(L371S+F375S)** | 4.81×10^5^ | 9.49×10^-2^ | 197 | 3.82×10^5^ | 1.53×10^-1^ | 400 | 6.31×10^5^ | 4.75×10^-2^ | 75.24 |
| **Omicron-(L371S+P373S+F375S)** | 7.27×10^5^ | 1.64×10^-3^ | 2.25 | 8.39×10^5^ | 1.63×10^-2^ | 19.39 | 7.32×10^5^ | 7.51×10^-3^ | 10.26 |
